# Evolution and multiple roles of the Pancrustacea specific transcription factor *zelda* in insects

**DOI:** 10.1101/126417

**Authors:** Lupis Ribeiro, Vitória Tobias-Santos, Danielle Santos, Felipe Antunes, Geórgia Feltran, Jackson de Souza Menezes, L. Aravind, Thiago Motta Venancio, Rodrigo Nunes da Fonseca

**Affiliations:** Laboratório Integrado de Bioquímica Hatisaburo Masuda, Núcleo em Ecologia e Desenvolvimento SócioAmbiental de Macaé (NUPEM), Campus UFRJ Macaé Av. São José do Barreto 764, ZIP CODE: 27965-550.; Laboratório de Química e Função de Proteínas e Peptídeos, Centro de Biociências e Biotecnologia, Universidade Estadual do Norte Fluminense Darcy Ribeiro, ZIP CODE: 21971-550; Instituto Nacional de Ciência e Tecnologia em Entomologia Molecular - INCT-EM; National Center for Biotechnology Information, National Library of Medicine, National Institutes of Health, Bethesda, MD 20894, USA.

## Abstract

Gene regulatory networks (GRN) evolve as a result of the coevolutionary process acting on transcription factors and the cis-regulatory modules (CRMs) they bind. The zinc-finger transcription factor (TF) *zelda* (*zld*) is essential for maternal zygotic transition (MZT) in *Drosophila melanogaster*, where it directly binds over thousand CRMs to regulate chromatin accessibility. *D. melanogaster* displays a long germ type of embryonic development, where all segments are simultaneously generated along the whole egg. However, it remains unclear if *zld* is also involved in MZT of short-germ insects (including those from basal lineages) or in other biological processes. Here we show that *zld* is an innovation of the Pancrustacea lineage, being absent in more distant arthropods (e.g. chelicerates) and other organisms. To better understand *zld’s* ancestral function, we thoroughly investigated its roles in a short-germ beetle, *Tribolium castaneum*, using molecular biology and computational approaches. Our results demonstrate roles for *zld* not only during the MZT, but also in posterior segmentation and patterning of imaginal disc derived structures. Further, we also demonstrate that *zld* is critical for posterior segmentation in the hemipteran *Rhodnius prolixus*, indicating this function predates the origin of holometabolous insects and was subsequently lost in long-germ insects. Our results unveil new roles of *zld* in maintaining pluripotent state of progenitor cells at the posterior region and suggest that changes in expression of *zld* (and probably other pioneer TFs) are critical in the evolution of insect GRNs.

## BACKGROUND

Gene regulatory networks (GRN) depend on the co-evolution of transcription factors (TFs) and their relationship with the cis-regulatory modules (CRMs) they bind to [1,2]. In insects, the detailed role of a number of TFs and CRMs have been well-described, particularly during the embryogenesis of the fruit fly *Drosophila melanogaster* [3].

In metazoans, the period following fertilization is typically characterized by rapid and near-synchronous mitotic divisions and cleavages, that occur under conditions of minimal cellular differentiation. Cleavages typically depend on maternally supplied factors and the zygotic genome transcription is constrained during this early period of development [4]. A conserved process of metazoan embryogenesis is the maternal-zygotic transition (MZT), which is characterized by two critical steps: 1) the elimination of a maternal set of mRNAs and proteins and; 2) the beginning of zygotic transcription, which leads to the zygotic genomic control of development [5]. In *D. melanogaster*, the majority of the first set of zygotic transcripts are regulated by a TF named *zelda (zld) [6]*, a zinc finger TF with particular affinity for promoter regions containing TAG Team sites [7–9] - heptamers constituted by CAGGTAG and its variants. *zld* (*Dm-zld*, in *D. melanogaster*) binding sites have been identified in *D. melanogaster* embryos (cycles 8 to 14) by chromatin immunoprecipitation coupled with high-throughput sequencing (ChIP-Seq) [7]. *Dm-zld* regulates a large set of genes involved in important processes such as cytoskeleton organization, cellularization, germ band development, pattern formation, sex determination and miRNA biogenesis [6]. *Dm-zld* was also suggested to participate in larval wing disc development [10] and its overexpression during wing imaginal disc formation led to wing blisters in adults, an indicative of improper adherence of ventral and dorsal wing epithelia [11]. Nevertheless, while *zld’s* functions have been thoroughly investigated in *D. melanogaster* MZT, its roles in other organisms and biological processes remain elusive.

*D. melanogaster* displays a long germ type of embryonic development, during which all segments are simultaneously generated along the whole egg. In contrast, short germ insects generate anterior (e.g. head) segments early in development, while the remaining segments are patterned from the posterior region, the growth zone (GZ). Since short germ development is considered to be the ancestral mode of development, short-germ insects have been established as developmental model systems [12,13]. The short-germ red flour beetle *Tribolium castaneum* (Tc) was the first beetle species to have its genome completely sequenced [14]. *T. castaneum* displays a short life-cycle and is amenable to gene silencing via RNAi [15], gene overexpression [16], specific tissue expression [17] and fluorescence labeling during early development [18]. Several developmental processes that have been investigated in *T. castaneum* were lost or extensively modified in the *D. melanogaster* lineage, such as GZ formation [19], extensive extra-embryonic morphogenesis [20] and the formation of a morphologically complex head during embryogenesis [21]. Early development of *T. castaneum* is similar to most other insect groups, in which synchronous rounds of division are followed by nuclei migration to the egg cortex and membrane segregation of nuclei into separate cells form the so called uniform blastoderm [18,22]. Taken together all the genetic and morphological information on *T. castaneum* early development, along with the established techniques mentioned above, *T. castaneum* is an ideal model to understand the evolution of *zld’s* function during insect development.

In the present work, we provide the first comprehensive analysis of *zld* orthologs across a wide range of species. Further, we provide functional analysis of a *zld* ortholog in a non-Drosophillid insect, the short germ beetle *T. castaneum* (*Tc-zld*). Among our main results are: 1) The identification of some previously overlooked conserved domains in Zld; 2) An inference of the evolutionary origin of *Zld*, based on phyletic analysis in various hexapods and crustaceans; 3) The identification of a conserved set of 141 putative *T. castaneum zld* (thenceforth referred as *Tc-zld*) targets (i.e. genes with upstream TAG team sequences), enriched in TFs, whose homologs that have been demonstrated to be *zld* targets in *D. melanogaster*; 4) Identification of key roles played by *Tc-zld* during the MZT; 5) Identification and experimental validation of two new biological roles of *zld* in *T. castaneum:* segment generation from the posterior GZ during embryogenesis and postembryonic imaginal disc development; 6) Demonstration that *zld* is also involved in GZ patterning in the hemipteran *Rhodnius prolixus*, supporting a conserved role in the GZ of a distant short-germ species. Altogether, our results unveil new roles of *zld* as a pioneer TF acting in various developmental processes across distantly-related insects.

## RESULTS AND DISCUSSION

### Identification of conserved domains and tracing the origin of *zld*

While previous studies reported that *zld* is involved in *D. melanogaster* MZT [6], its evolutionary history remains unclear. We investigated the phyletic distribution of *zld* and found a single ortholog in all the inspected insect genomes, including the beetle *T. castaneum* (Figure 1; Table S1), indicating that insects are sensitive to increased copy number of this important regulator, which is interesting considering that different TF families are particularly prone to expansions across insect lineages [23]. Further, *zld* homologs were also found in some (but not all) collembolan and crustacean genomes (Figure 1; Table S1). The canonical architecture of Zld has been reported as comprising a JAZ zinc finger (Pfam: zf-C2H2_JAZ) domain and a C-terminal cluster of four DNA binding C2H2 zinc finger domains (zf-C2H2) [6,11,24]. However, we performed a sensitive and detailed analysis of Zld proteins from multiple species and found other notable conserved protein domains and structural features. Firstly, we found two additional zf-C2H2 domains, N-terminal to the JAZ domain (Figure 1). The most N-terminal zf-C2H2 is absent or partially eroded (i.e. without the conserved cysteines and histidines) in some species (Figure 1), including *D. melanogaster*. This observation was confirmed by checking *Dm-zld* alternative splicing isoforms (data not shown). On the other hand, the second N-terminal zf-C2H2 domain is conserved in virtually all extant insects, but absent or degenerate in the other Pancrustaceans, (e.g. *Daphnia magna*, it is partially conserved except the loss of one cysteine) (Figure 1). Further, between this second zf-C2H2 and the JAZ domain, there is a strikingly conserved acidic patch, which is likely to be involved in the recruitment of chromatin remodeling proteins based on the conservation of a similar acidic patch in the chromatin remodeler CECR2 [25] (Figure 1). This patch is characterized by an absolutely conserved motif of the form [DE]I[LW]DLD which is likely to adopt a helical conformation amidst surrounding disordered regions. Analogous acidic patches are also seen in the HUN domain histones chaperones [26] A largely disordered region, between the JAZ finger and the cluster of 4 widely-conserved zf-C2H2 domains, has been shown to be important for *Dm-zld* transactivation *in vitro* [24]. Our analysis of evolutionary constraints on the protein sequence revealed a motif of the form hP[IVM]SxHHHPxRD which appears to be under selection for retention despite the strong divergence in this region. Hence, it is possible that it specifically plays a role in the observed role in transactivation. Further, between the two N-terminal zf-C2H2 domains, there is a conserved motif, RYHPY, that is highly conserved in Zld proteins and could be involved in nuclear localization (Figure 1), as predicted for other transcriptional repressors [27]. Given the conservation of these additional domains, we hypothesize that they also play important roles in Zld functions that were previously attributed exclusively to the C-terminal domains.

**Figure 1:**
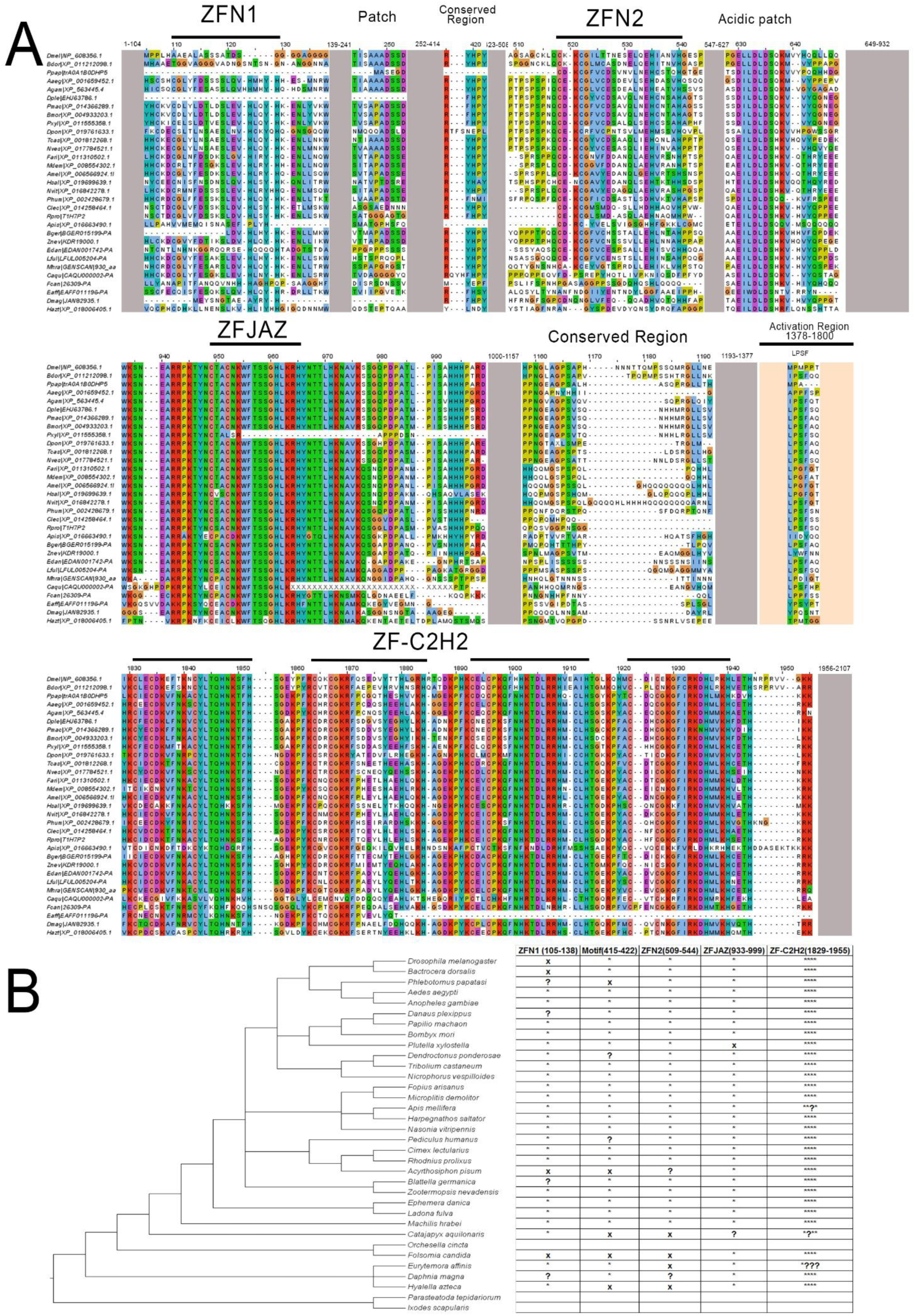
*Tc-zelda* proteins from insects and crustaceans. (A) Multiple sequence alignment of zelda proteins, representing major groups of arthropods. (B) Conserved protein architecture features of Zelda proteins. Asterisks and x marks represent presence and absence, respectively. Question marks denote that the feature is either partially preserved or could be flagged as absent due to sequencing or assembly errors (e.g. wrong start codons). The cladogram was organized according to a previously reported phylogenetic study [67]. Two outgroups without *zld* orthologs were also included.

Aiming to elucidate the origins of *zld*, we performed extensive sequence searches and were unable to find homologs with the conserved domain architecture outside of the Pancrustacea, indicating that *zld* is an innovation of this lineage. We detected clear *zld* homologs across insects, including the termite *Zootermopsis nevadensis* (Order Isoptera), the scarce chaser *Ladona fulva* (Order Odonata), the mayfly *Ephemera danica* (Order Ephemeroptera) and *Machilis hrabei* (Order Archaeognatha). *zld* homologs were also found in crustaceans belonging to different classes (*Daphnia magna, Hyalella azteca* and *Eurytemora affinis*), as well as in the collembola *Folsomia candida*. Curiously, we found no *zld* in the genome of *Orchesella cincta* (collembola) and *Daphnia pulex* (crustacean), suggesting either that it is not absolutely conserved outside of insects or is missed due to incompleteness of the deposited genomes. Specifically, we carefully searched the genomes of other non-insect arthropods, including chelicerates (e.g. the tick *Ixodes scapularis* and the spider *Parasteatoda tepidariorum*), and found no proteins with the canonical Zld domain organization (Figure 1). General searches on Genbank against non-Pancrustacea arthropod proteins also returned no Zld orthologs. Although BLAST searches with Zld proteins against the *nr* and *refseq* databases recover a number of significant hits in several distant eukaryotes, the similarity is almost always restricted to the C-terminal cluster of four Zf-C2H2 domains, which are very common across many TF families (e.g. *glass, earmuff/fez, senseless/gfi-1 and jim*). Taken together, our results support the early emergence of *zld* in the Pancrustacea lineage, with possible subsequent losses in particular species. Importantly, all the insect genomes we inspected have exactly one *zld* gene, indicating that this gene became essential in hexapods.

### *Tc-zld* is a master regulator of signaling genes and other transcription factors

Previous studies in *D. melanogaster* using microarrays and ChIP-Seq revealed that *Dm-zld* regulates the transcription of hundreds of genes during early embryogenesis [6,7,28]. In *D. melanogaster*, enhancers bound by Dm-Zld are characterized by a consensus sequence CAGGTAG (i.e. TAG team sequence), which is overrepresented in early zygotic activated genes, including TFs involved in AP and DV patterning [8]. Since the TAG team motif identified in *D. melanogaster* is conserved in *A. aegypti* [29], we investigated whether we could predict *Tc-zld* targets by detecting upstream TAG team motif.

Firstly, an *ab initio* approach using DREME [30] was employed to analyze 2kb upstream regions of all *T. castaneum* protein-coding genes. This analysis uncovered a motif (i.e. GTAGGTAY) that is nearly identical to the TAG team motif (Figure 2A).

**Figure 2:**
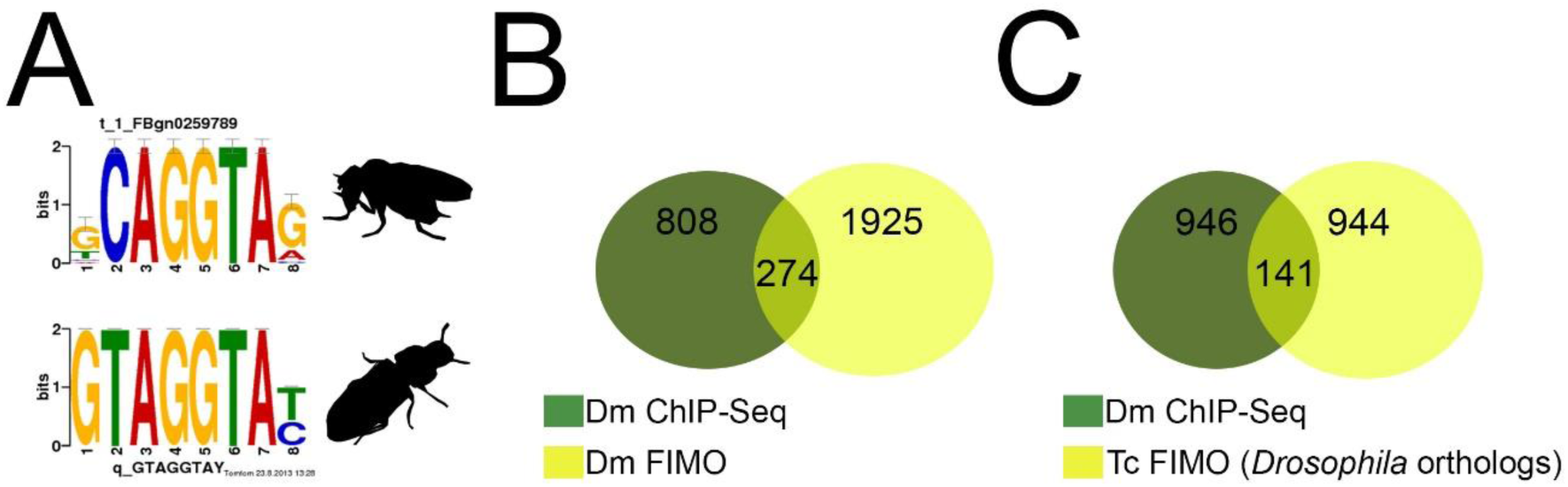
Computational identification of *Tc-zelda* target genes. (A) TOMTOM comparison of *D. melanogaster* motif similar to the TAG team [8] obtained by DREME and putative *Tribolium castaneum zld* DREME motif. (B) Venn diagram of the *D. melanogaster* ChIP-Seq MZT regulated genes (green) from [7] and *D. melanogaster* genes predicted by FIMO analysis with the putative DREME motif (yellow). (C) Venn diagram of the *D. melanogaster* Dm-Zld MZT targets [7] (green) and *D. melanogaster* orthologs (only one-to-one) of putative *Tc-zld* targets.

We used the *D. melanogaster* genome and experimental data [6,7,28] to validate our approach and found a significant overlap between experimental and predicted *zld* targets in *D. melanogaster* (Figure 2B). The putative *T. castaneum* motif was then used to screen the *T. castaneum* genome, resulting in the identification of 3,250 putative *zld* targets, representing ∼19% of the *T. castaneum* genome (Figure 2C, Table S2). Comparison of the putative *Tc-zld* targets with 1,087 genes regulated by Zld during *D. melanogaster* MZT [7,28] allowed the identification 141 *D. melanogaster* genes for which one-to-one orthologs figured among the putative *Tc-zld* targets (hypergeometric distribution, *P* < 4.5 × 10^−4^; Figure 2C, Table S3). Functional analysis of this gene set using DAVID [31] uncovered the enrichment of important categories, including a highly significant cluster of 26 homeobox TFs (Table S4) and other significant clusters comprising genes involved in regionalization and segment specification, imaginal disc formation and metamorphosis (Table S4). Interestingly, this gene set included multiple developmental regulators such as anteroposterior (AP), gap, pair-rule, homeotic and dorsoventral (DV) genes (Table S2). Since several of these genes are involved in early developmental processes, we focused our initial analysis of *Tc-zld’s* function during embryogenesis.

### *Tc-zld* expression is progressively confined to the posterior region during growth-zone formation

Since *zld* is maternally expressed at the germ line in *D. melanogaster* [6], we compared its transcription in *T. castaneum* female ovaries and carcass by RT-qPCR. *T. castaneum* early development starts with synchronous divisions during the first three hours of embryogenesis (at 30°C), followed by nuclei migration to the egg cortex and membrane segregation of nuclei into separate cells, approximately 7-8 hours after egg lay [18,22]. We found that *Tc-zld* is highly expressed in the ovaries (Figure 3A), supporting the transcription of *Tc-zld* in the germ line. The abundance of *Tc-zld* transcripts is also higher in the first three hours of development than in the next two 3-hour periods (i.e. 3-6 and 6-9 hours) (Figure 3B), suggesting that *Tc-zld* mRNA is degraded after the first 3 hours of development. An antibody against the transcriptionally active form of RNA polymerase II, previously used in other arthropod species [32,33], showed that zygotic transcription in *T. castaneum* begins between three and six hours of development, shortly after the nuclei have reached the periphery (Figure S1). *In situ* hybridization confirmed maternal expression of *Tc-zld* in the first three hours of development (Figure 3C,D) and showed a progressive confinement of *Tc-zld* mRNA to the posterior region of the egg, where the germ rudiment will be formed (Figure 3E,F). Between 6 and 9 hours of development, *Tc-zld* expression is further restricted to the posterior region (Figure 3G,H), where the GZ will generate new segments [19,34]. Later in development, *Tc-zld* expression is still detected at the GZ and nervous system (Figure S2). Although *zld* is expressed at the neural progenitors in *D. melanogaster* [6,11] and *T. castaneum*, biased posterior expression at the GZ is a specific feature of short-germ insects like *T. castaneum*.

**Figure 3:**
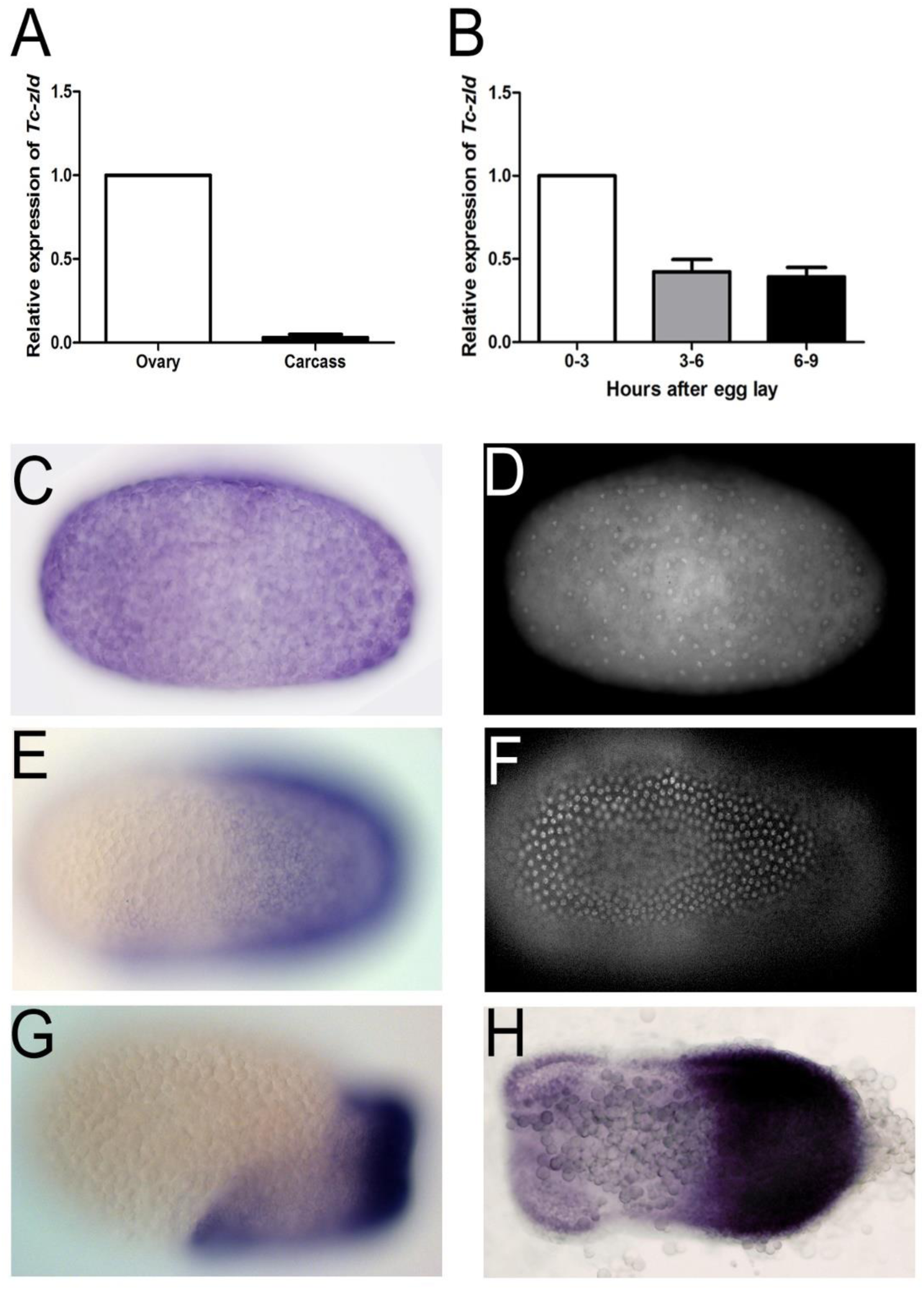
*Tc-zelda* is maternally provided and progressively confined to the posterior growth-zone during embryonic development. (A) Relative expression of *Tc-zld* in ovary and carcass. (B) Relative expression of *Tc-zld* 0-3, 3-6 and 6-9 hours after egg lay. Expression was normalized in relation to the constitutive gene rps3 in both experiments. (C) Pre-blastoderm stage embryo (0-3 hours) shows *Tc-zld* transcripts uniformly distributed and its respective DAPI staining in D. (E) At uniform blastoderm stage (3-6 hours) transcripts begin to occupy the germ rudiment, respective DAPI in F. (G) Shortly before posterior invagination *zld* expression starts to concentrate at the posterior region where growth zone will form (6-9 hours). (H) Shortly before the beginning of germ band elongation *zld* expression is largely confined to the GZ.

### Parental injection of *Tc-zld* dsRNA reduces number of eggs laid and impairs embryogenesis

It has been shown that injection of dsRNA in *T. castaneum* females, parental RNAi (pRNAi) reduces maternal and zygotic mRNA expression of a given gene [15]. We injected *zld* dsRNA in females and analyzed its transcriptional levels during embryogenesis by qRT-PCR. After *zld* pRNAi, *Tc-zld* mRNA levels were reduced in the first two weeks of egg laying, severely impairing larval hatching (Figure 4A-C).

**Figure 4:**
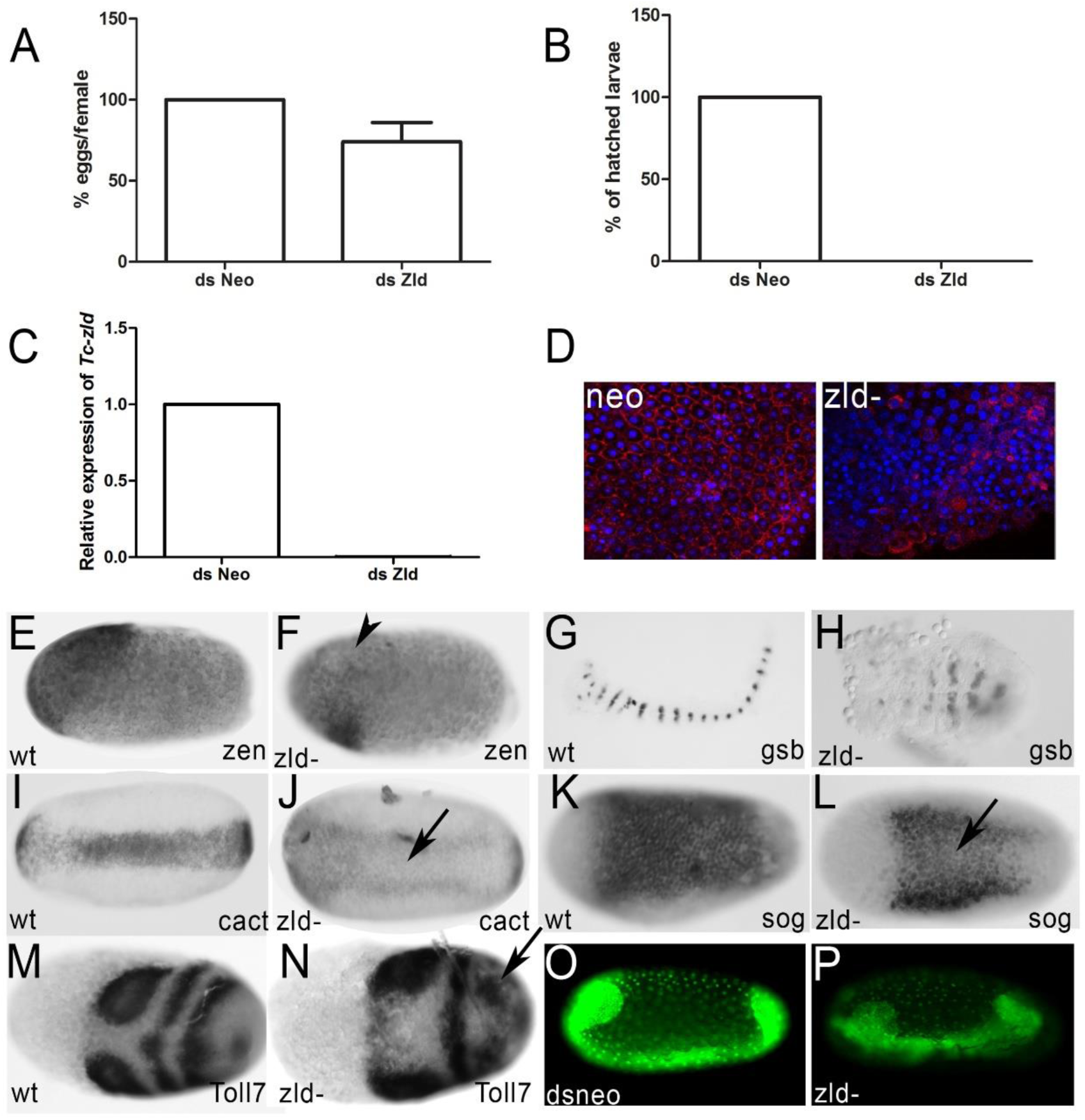
*Tc-zelda* parental RNAi affects oviposition and cellularization while embryonic RNAi affects growth zone patterning. (A) Normalized number of *dsneo* and *zld* RNAi collected eggs. *zld* pRNAi reduces oviposition. (B) Percentage of hatched larvae obtained from eggs after *zld* pRNAi in comparison to the control (*dsneo* RNAi). *zld* pRNAi leads to lethality during embryogenesis. (C) Relative expression of *zld* in *dsneo* RNAi (control) and *dszld* RNAi during the first two egg lays. *zld* pRNAi almost completely abolishes *zld* expression. (D) Phospho-tyrosine (red) and DAPI (blue) staining in *dsneoand zld* RNAi eggs during cellularization. *zld* pRNAi impairs cellularization. (E,F) *Tc-zen* expression in the serosa [35] is reduced after *Tc-zld* pRNAi (arrowhead) when compared to the WT. (G,H) Segment generation from the growth-zone (GZ) is disrupted, as judged by the analysis of *Tc-gsb* expression after *Tc-zld* pRNAi (H) and *dsneo* (G). Analysis of other markers is provided at Figure S4. (I,J) The expression of dorsoventral gene *Tc-cactus* [37] was reduced after *Tc-zld* pRNAi (J-arrow) when compared to the (I) control. (K,L) *Tc-zld* dsRNA embryonic injection (eRNAi) into nuclear GFP transgenic line (Sarrazin et al., 2012) affects segment generation from the posterior GZ (L-asterisk), when compared to nGFP embryo injected with *dsneo* dsRNA

Importantly, identical knockdown phenotypes during embryogenesis were obtained by using a second, non-overlapping, dsRNA construct (Figure S3). Further, morphological analyses showed that cellularization was severely disrupted in over 50% of the *zld* dsRNA embryos (Figure 4D), similarly to previously reported in *D. melanogaster zld* mutants [6,11]. The remaining *zld* pRNAi embryos were not severely affected during cellularization and developed beyond that stage. Finally, we also found that putative conserved target genes were down-regulated in embryos after *zld* pRNAi, such as the early zygotic genes involved in AP patterning, the serosal gene *Tc-zerknullt* [35] (Figure 4E,F) and the gap gene *milli-pattes* [36] (Figure S4).

Changes in the spatial distribution of transcripts from predicted dorsoventral target genes were also observed after *Tc-zld* RNAi. In WT embryos, the TF Dorsal forms a dynamic transient gradient, which activates *Tc-cactus* (*Tc-cact*) and *Tc-short-gastrulation* (*Tc-sog*) at the ventral region [22,37]. After *Tc-zld* pRNAi, the expression of *Tc-cact* and *Tc-sog* is observed in two lateral domains, in contrast to the expression at the single ventral domain in dsneo embryos (Figure 4I,J,K,L-arrow). These results suggest that *Tc-zld* is required for proper activity of Dorsal at the ventral-most region of the embryo.

As discussed above, *Tc-zld* is highly expressed at the posterior region of the embryo (Figure 3) and likely associated with segmentation and regionalization (Table S4). Further, several putative *Tc-zld* targets are involved in posterior segmentation, such as *caudal (Cdx), even-skipped (Eve)*, and several Hox genes such as *Ultrabithorax, Abdominal-A* and *Abdominal-B*. *Tc-eve*, for example, is essential for the establishment of a genetic circuit required for posterior segmentation [38]. Interestingly, *Tc-zld* pRNAi embryos showed a continuous *Tc-eve* expression domain instead of the typical stripe patterning required for WT segmentation (Figure S4).

Elegant studies on *T. castaneum* GZ patterning showed that cell proliferation is not essential for segment generation, which rather occurs by coordinated cell movement and intercalation [19,39]. We then evaluated whether *Tc-zld* regulates genes involved in cell intercalation, such as *Toll2, Toll6* and *Toll8* [40,41]. Interestingly, *Toll7* (TC004474) and *Toll8 (Tollo*:TC004898) are among the common *zld* targets conserved between *D. melanogaster* and *T. castaneum*. Since *Tc-Toll7* is expressed during early segmentation [41], we compared its expression in *dsneo* and *Tc-zld* RNAi embryos. While anterior expression of *Tc-Toll7* is not affected, the striped expression at the posterior region is lost when *Tc-zld* expression is reduced (Figure 4M,N). In summary, *Tc-zld* regulates the expression of several genes that are critical for early AP (*zen, mlpt*) and DV (*sog, cact, twist*) patterning and, in a second phase, genes required for posterior elongation (e.g. *Toll7*).

### *Tc-zld* plays specific roles in the posterior GZ

While pRNAi diminishes maternal and zygotic expression of *Tc-zld* in *T. castaneum [15]*, embryonic dsRNA injections (eRNAi) may affect only the zygotic component [37,42]. To investigate if *zld* is specifically required for embryonic posterior patterning, we injected *Tc-zld* dsRNA in transgenic embryos expressing nuclear GFP (nGFP), as previously described [18,19]. Embryonic injections of *zld* dsRNA after MZT (see methods for details) impaired segment generation from the GZ, while *dsneo* injected embryos developed like WT ones (Figure 4G,H, Sup. movies 1,2). In addition, expression of the predicted target *Tc-eve*, a key TF involved in GZ patterning [34] has been largely down-regulated upon *zld* embryonic RNAi (data not shown), as previously observed for *zld* pRNAi (Figure S4). In summary, our results imply that *Tc-zld* is involved not only in the MZT, early patterning and nervous system formation, as described for *D. melanogaster* [6], but also play roles in segment generation from the GZ, a structure found only in embryos of short-germ insects like *T. castaneum*.

### Functional analysis in the hemipteran *Rhodnius prolixus* shows a conserved role of *zld* in short-germ insect posterior region

Although we demonstrated the involvement of *zld* in *T. castaneum* GZ, the conservation of this regulatory mechanism in other species remained unclear. Thus, we sought to analyze the functions of the *zld* ortholog in the hemipteran *R. prolixus* (*Rp*), which is a hemimetabolous insect and lacks the complete metamorphosis present in holometabolous species such as *T. castaneum* [43,44]. *Rp-zld* knockdown via pRNAi resulted in two types of embryonic phenotypes: 1) severe defects in gastrulation and lack of any appendage development (data not shown); 2) embryos which only developed the anterior-most embryonic regions comprising the head, gnathal and thoracic segments (Figure S5). Thus, *zld* is involved in posterior patterning not only in the beetle *T. castaneum*, but also in other short-germ basal insects, such as *R. prolixus*. To our knowledge this is the first direct description of *zld* function in insects other than *D. melanogaster*. Finally, a recent report showed *zld* as maternally transcribed in the hymenoptera *Apis mellifera* (also a long-germ insect), while zygotic transcripts are concentrated at the central region of the embryo during blastodermal stages [45].

### *Tc-zld* is essential for patterning of imaginal disc-derived structures: wing, legs, elytra and antenna

DAVID analysis of *D. melanogaster* orthologs of the putative *Tc-zld* targets uncovered a cluster of 29 genes involved in imaginal disc development (Table S4; GO:0007444). Among these genes are several homeodomain TFs, such as *distalless (Dll), Abdominal A (Abd-A), Abdominal B (Abd-B), zen, Engrailed (En), caudal (cad), defective proventriculus (dve), mirror (mirr), araucan (ara-iroquois), Drop (dr)* other TF such as *dachsund (dac), taranis (tara) and Lim1, PoxN, kn, sob, drm, Awh, dp*.

As the first step towards the characterization of the post-embryonic role of *zld*, we analysed its expression by qRT-PCR in larvae of third (L3), fifth (L5) and seventh (L7) stages, and first pupal stage (P1) (Figure 5A-D). Interestingly, *Tc-zld* expression increases during successive larval stages and sharply decreases after pupal metamorphosis (Figure 5E). This suggests that *Tc-zld* might be required for late larval stages taking place during larvae-pupae metamorphosis, such as growth and patterning of structures derived from imaginal discs in *D. melanogaster*, such as antenna, legs, fore and hindwing [46,47]. Further, we found that three out of five predicted *Tc-zld* targets, namely *Dll, Wg* and *Lim-1*, also displayed an increase in expression during late larval and pupal development (Figure 5E).

**Figure 5:**
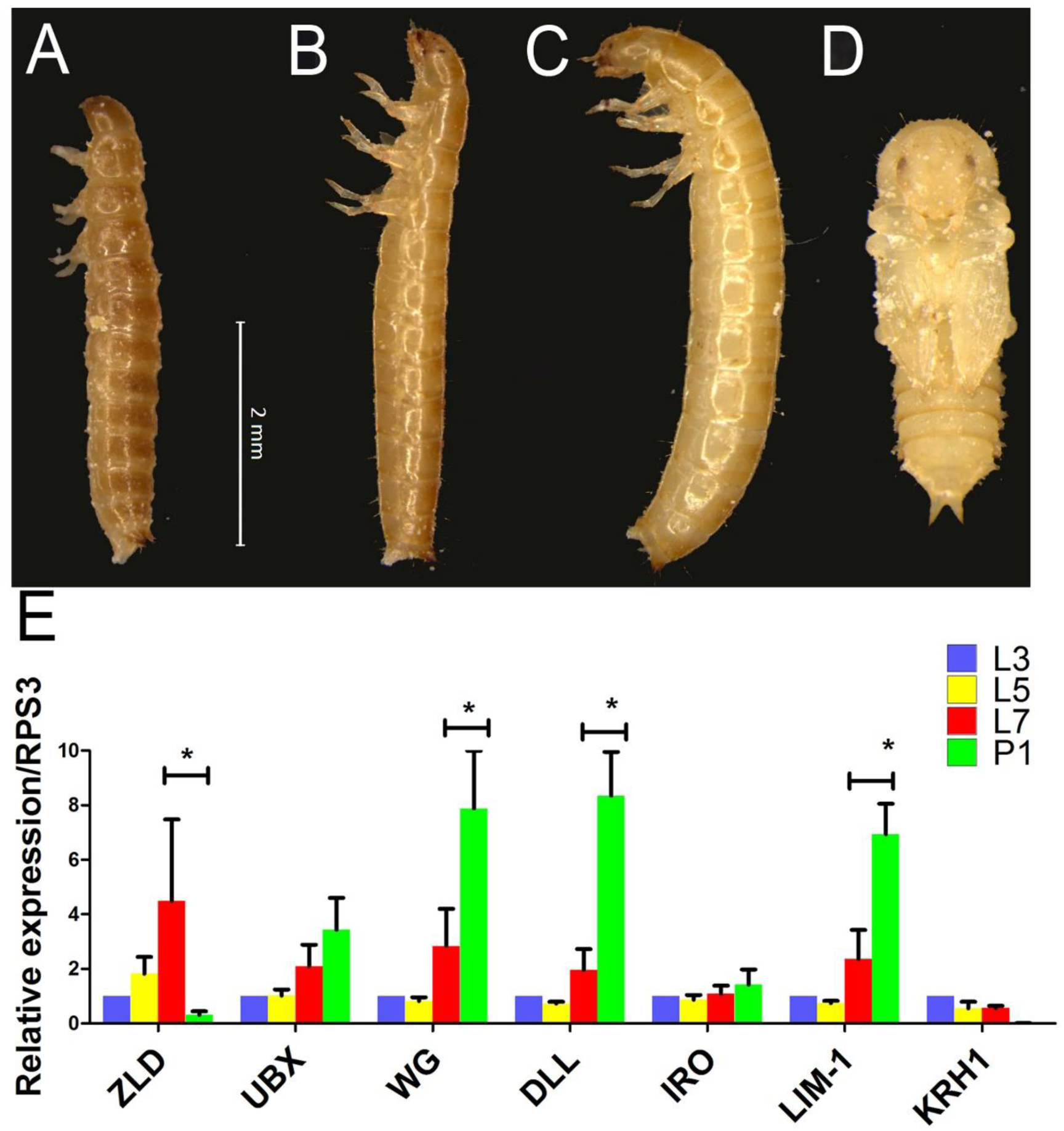
Larval stages and expression dynamics of *zld* and its putative target genes. (A-D) Morphology of *Tribolium castaneum* larvae on 3rd (A-L3), 5th (B-L5), 7th (C-L7) and early pupal stages (D-P1). (E) Relative expression of *zld, Ubx, wingless (wg), distalless (dll), Iroquois (iro), Lim-1* and *Kruppel-homolog-1 (KRH-1)* at L3, L5, L7 and P1. Asterisks represent significant differences between stages (P<0.05).

To investigate *zld’s* post-embryonic roles, we injected two non-overlapping *Tc-zld* dsRNA constructs into early (L3) and late larval (L6) stages, as previously described [47]. qRT-PCR confirmed that *Tc-zld* was down regulated after dsRNA injection (Figure 6A). *Tc-zld* dsRNA injections at early larval stages (L3) led to over 50% lethality during pupal stages (data not shown). Atypical adult pigmentation in the pupal head and reduction in the wing size were observed after early *Tc-zld* dsRNA injection, suggesting that proper *Tc-zld* expression is required for metamorphosis and wing growth (Figure 6B,C). Interestingly, *Tc-zld* dsRNA injections at late larval stage (L6) displayed a different phenotype when compared with early larval dsRNA injections (L3). L6 dsRNA injected larvae reach adulthood in comparable numbers to *dsneo* injected larvae (Figure 6D). Specifically, *Tc-zld* dsRNA adults showed a series of morphological alterations in tissues undergoing extensive morphological changes during metamorphosis, such as fore and hind wings, antennae and legs (Figure 6E,F, 7 and 8).

**Figure 6:**
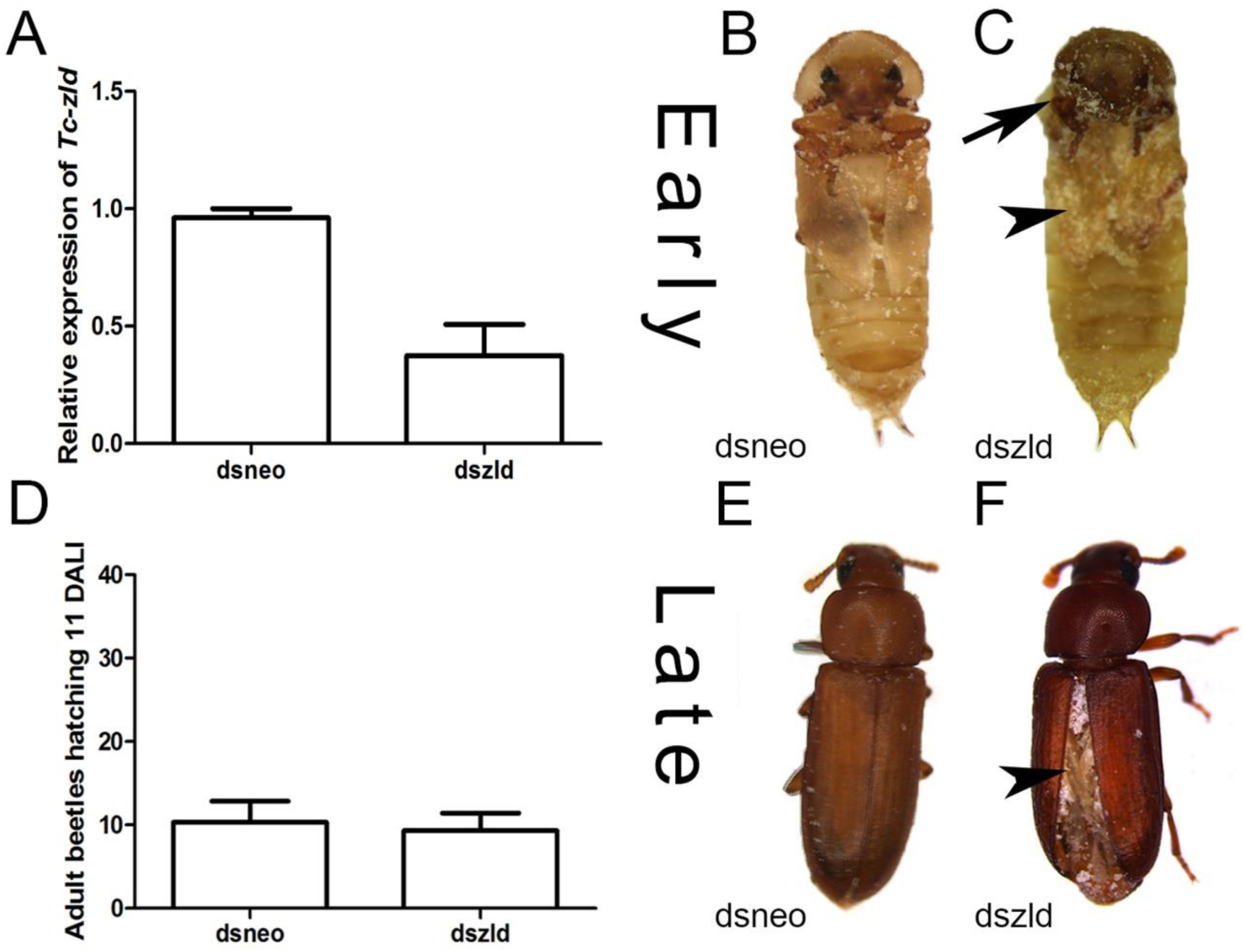
*Tc-zelda* larval RNAi affects elytra enclosure and molting. *Tribolium castaneum* larvae were injected during early (3rd) and late (6th) stages as previously described [47,68]. (A) Relative expression of *Tc-zld* during pupal stages after *Tc-zld* dsRNA or *neo* dsRNA. qRT-PCR was normalized using *Tc-rps3* gene, as previously described [66]. (B,C) Morphology of late pupae obtained from larvae injected with *zld* or *neo* dsRNA. Differential pigmentation in the head (arrow) and reduced wings (arrowhead) were observed in *zld* dsRNA and not in *dsneo* dsRNA pupae. (D) Number of emerging adult beetles eleven days after larval injection (DALI). (E,F). Adults obtained by late larval injections of *neo* (E) or *zld* (F) dsRNA. Hindwings are usually not properly folded underneath the forewing (elytra).

**Figure 7:**
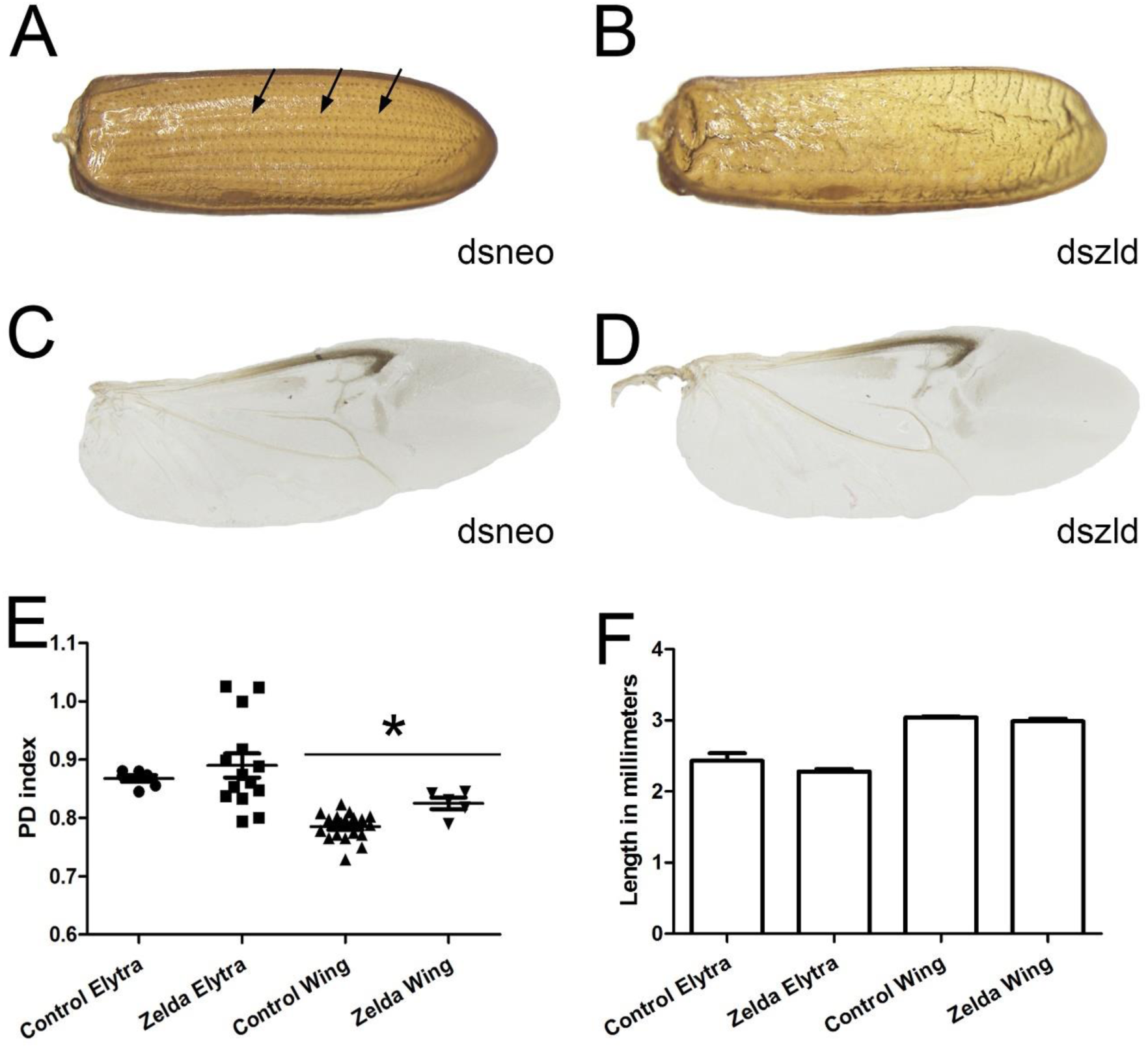
*Tc-zld* knockdown in larval stages affects elytra and wings in adult stage: (A) Control elytron extracted from a *dsneo* adult shows a parallel venation pattern and rigid chitinous structure. (B) *Tc-zld* RNAi elytron displays a disrupted vein pattern and a less resistant structure. (C,D) Hindwings were dissected and photographed. Overall morphological pattern is not affected in both wings, although *zld* dsRNA wings show severe dehydration after ethanol fixation (data not shown). (E) PD indices comparison between *dsneo* and *dszld* hindwings and forewings, a PD index reflects the shape of the wing based on its dimensions ratio [49]. Statistical analysis carried out by unpaired t-test assuming unequal variances (*=P<0.0001, ns= no significance) indicating that *Tc-zelda* silencing affects hindwing shape. (F) *Tc-zld* RNAi elytra and hindwing does not show statistically significant differences in length compared to respective *neo*.

**Figure 8:**
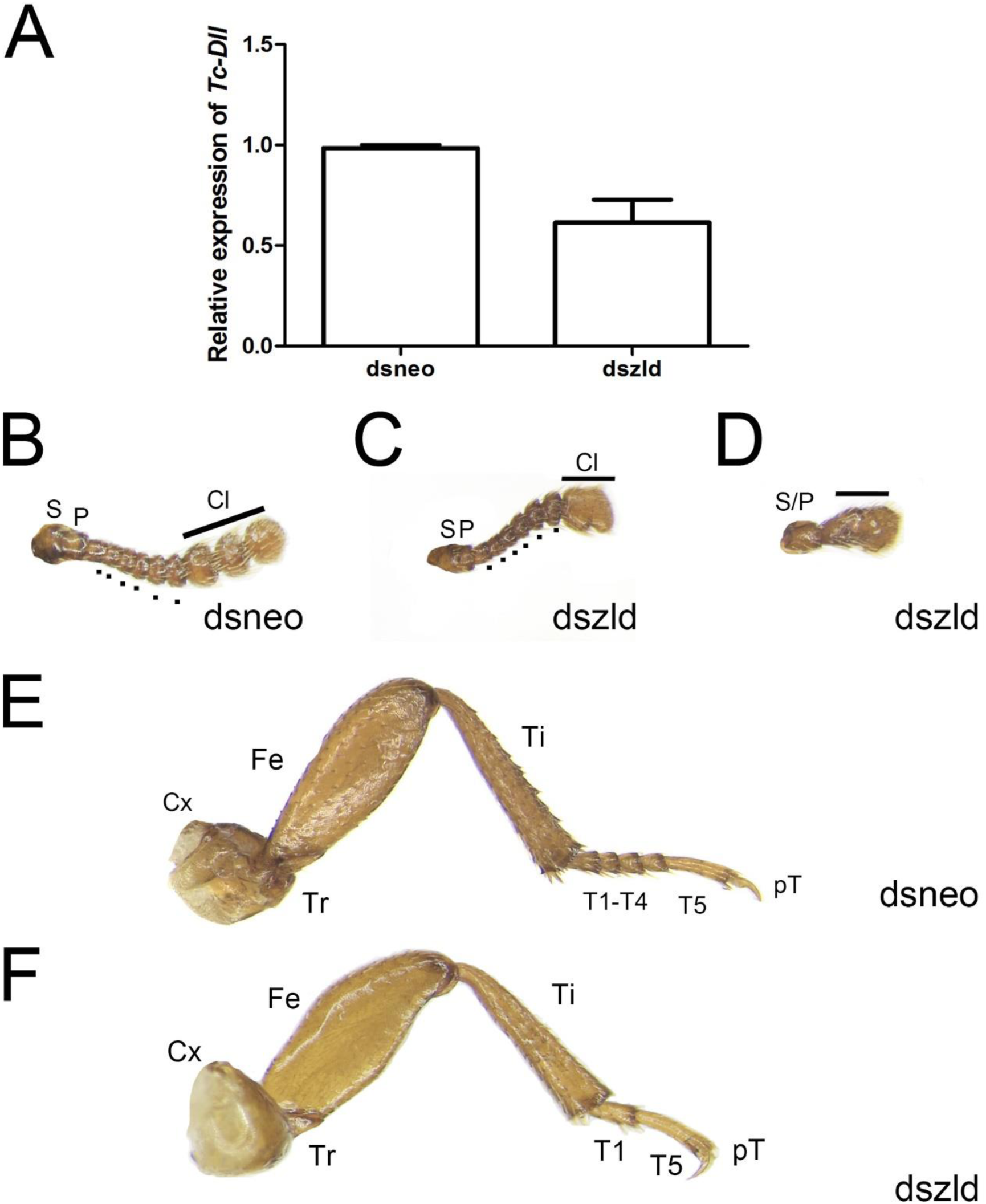
*Tc-zld* knockdown induces phenotypes in antennae and legs of adult *T.castaneum*: (A) Relative expression of *Tc-Dll* at pupal stages after dsneo or dszld injections. (B-D) Antennae extracted from control injected individuals display 11 segments, including two proximal (S= scape, P = pedicel), six intermediate (Flagellum=dots) and three distal modified segments that form the club (Cl). (E,F) Injection of *dszld* caused caused fusion club segments while in stronger phenotypes the antennae fail to segment, developing as an antennal rudiment. Legs of *T.castaneum* have six podomeres: Coxa (Cx), Trochanter (Tr), Femur (Fe), Tibia (Ti), Tarsus (T) and pretarsus (pT). (D) mesothoracic legs of *dsneo* injected individuals five tarsal segments, or tarsomeres. (E) In *Tc-zld* silenced individuals suffer some tarsal segments are deleted resulting in a reduced tarsus, while other leg structures show no defects.

The most visible effect of *Tc-zld* dsRNA beetles was a failure of the forewings (elytra) to enclose the hindwings, leading to the exposure of the dorsal abdomen (Figure 6E,F). Elytra, which are highly modified beetle forewings, have been proposed as an important beetle innovation, being required for protection against mechanical stresses, dehydration, and predation [48]. In line with this hypothesis, *Tc-zld* dsRNA adults with exposed abdomens started to die a few days after metamorphosis, probably due to dehydration.

Next, we performed a detailed morphological analysis to investigate if patterning defects resulting from *Tc-zld* dsRNA occurred in the sclerotized elytra (forewing) or in the hindwing (Figure 7A-D). This analysis showed that the parallel vein pattern of the elytra (arrows, Figure 7A) is disrupted after *Tc-zld* dsRNA in comparison to the control (Figure 7A,B). Nevertheless, hindwings of *Tc-zld* dsRNA beetles showed no signs of abnormal venation (Figure 7C,D). To address if fore or hind wing shapes have changed upon *Tc-zld* dsRNA knockdown, we applied the recently developed proximo-distal (PD) index [49]. While the PD index values of forewing (elytra) of *Tc-zld* and controls were similar, a slight but significant increase in PD index of the hindwings was observed (Figure 7E). Neither fore-nor hindwings length was altered upon *Tc-zld* knock down (Figure 7F). In conclusion, *Tc-zld* is required for proper venation pattern, but not shape, of the elytra. Previous analysis of *zld* expression in *D. melanogaster* showed expression in wing imaginal discs, particularly where mitotically active cells are located [11]. Moreover, *Dm-zld* overexpression during wing imaginal disc formation leads to adult wing blisters or tissue loss [10,11]. Nevertheless, to our knowledge, our study provides the first direct evidence of *zld’s* role in insect wing formation.

Interestingly, all *Tc-zld* dsRNA beetles that showed this ‘opened wing’ phenotypes (Figure 6F) also displayed defects in the legs and antennae. Antennae and legs share similar developmental gene networks, the so called serial homologs. *Distaless (Dll)*, one of the putative *Tc-zld* targets, is essential for appendage segmentation [50]. *Tc-Dll* is also expressed during late embryogenesis on the distal part of the leg and, as its name suggests, disruption of its function leads to the absence of distal leg and antennae segmentation [51]. Interestingly, we found that *Tc-zld* RNAi resulted in a significant decrease of *Tc-Dll* mRNA levels (Figure 8A), indicating that this gene is indeed downstream of *Tc-zld*.

Insect antennae possess three primary segments: scape, pedicel and flagellum. In *T. castaneum*, the adult antennae display eleven segments, out of which nine form the flagellum. The three most distal flagellar segments are enlarged and form the club, while the six intermediate flagella are called the funicle (Figure 8B) [52]. In mild phenotypes, *dszld* caused a joint malformation in distalmost flagellar segments resulting in fusion of the club (Figure 8C). On the other hand, strong *Tc-zld* phenotypes resulted in fusion of the scape and pedicel, leading to severe loss of flagellar joints and formation of a single truncated segment (Figure 8D). In contrast, no differences in scape and pedicel were observed.

*T. castaneum* legs originate during late embryogenesis and can be recognized as a small outgrowth of the body wall, the limb bud. In the adult stage, there are three pairs of segmented legs with six segments: coxa, femur, trochanter, tibia, tarsus and pretarsus [53]. In *T. castaneum* the tarsus is subdivided into smaller segments (i.e. the tarsomeres), five in prothoracic and mesothoracic legs and four in metathoracic legs. This tarsal segmentation occurs during beetle metamorphosis and this subdivision of the tarsus evolved in the common ancestor of insects, since the tarsus is not subdivided in non-insect hexapods [54]. After *Tc-zld* RNAi tarsal segments were absent or fused, resulting in leg shortening; in *dsneo* insects, legs were identical to that of wild-type animals (Figure 8 E,F). This indicates that some *Tc-zld* targets are involved in segment development or joint formation. Interestingly, a large domain of *Dll* expression is observed in beetle leg tarsus [55], and *Tc-Dll* knockdown in beetles also generated legs with tarsomere deletion [55,56].

### *zld* is a pioneer regulator of chromatin status during early embryogenesis and postembryonic development

Besides the conservation of *zld* roles in the MZT, our results uncovered two new biological roles of *zld* in the beetle *T. castaneum:* 1) regulation of segment generation from the posterior GZ during embryogenesis and; patterning of imaginal disc derived structures. But what do posterior GZ patterning, MZT and imaginal disc development have in common? All these processes involve extensive morphological changes and modifications of the differentiation status of precursor cells.

During *D. melanogaster* MZT, *zld* acts as a pioneer TF marking the chromatin of earliest expressed genes [7,57]. *Dm-zld* increases chromatin accessibility of the most important TFs involved in DV, dorsal, and AP patterning, *bicoid* [58,59]. Addition or removal of *Dm-zld* binding sites influences the timing of activation of Dorsal early zygotic targets in *D. melanogaster* [28,59], suggesting that Zld acts as a developmental timer. Our results showed an extensive maternal contribution of *Tc-zld* mRNAs, coupled with a zygotic *Tc-zld* expression in an anteroposterior progression (Figure 3). Shortly before gastrulation, *Tc-zld* is largely confined to the posterior region. We propose that *Tc-zld* mediates a progressive anteroposterior opening of the chromatin in *T. castaneum*, while posterior GZ cells retain an undifferentiated state. In zebrafish, Pou5f1, a homolog of the mammalian pluripotency TF Oct4, occupies SOX-POU binding sites before the onset of zygotic transcription and activates the earliest zygotic genes [60]. Moreover, Pou5f1 and Sox2 binding sites, chromatin states, and Pol II binding are similar in zebrafish embryos and mammalian ES cells, suggesting that the ancestral function of the vertebrate pluripotency factors were the activation of zygotic genes and control the developmental timing during early embryogenesis. We propose that posterior cells expressing *zld* maintain a pluripotent state during early stages of germ band elongation, while undergoing convergent extension movements [19,39]. Loss of *Tc-zld* expression might lead to premature tissue differentiation, lack of convergent extension and, ultimately, segmentation failure (Figure 4). Since *zld* is also expressed and important for posterior region of the hemimetabolous insect *R. prolixus* (Figure S5), *zld’s* role in posterior region dates back at least to the last common ancestor of Paraneoptera. *Tc-zld* might also play a similar role in the regulation of leg and antenna segmentation during metamorphosis (Figure 7, 8). The number of tarsomere segments in the leg and intermediate funicle are reduced after *Tc-zld* dsRNA injection, suggesting that the post-embryonic role of *Tc-zld* might be the accurate generation of the precursor cells required for appendage segmentation, as observed in the posterior embryonic region.

The evolutionary success of hexapods is attributed to a combination of features: their segmented body plan and jointed appendages, which were inherited from their arthropod ancestor and; wings and holometaboly, two features that arose later (Nicholson et al. 2014). It is interesting to notice that the specific Pancrustacea gene *zld* is required for most of these processes such as embryonic segment formation, wing (elytra) patterning, and appendage (antennae and leg) formation during beetle development. Like several other Zn-finger proteins which show tremendous lineage-specific diversity in eukaryotes, *zld* appears to have specific arisen within Pancrustacea and risen to play an important role as a “master TF”. Hence, future studies to understand how this newly emergent transcription factor was integrated into conserved backdrop of development-regulating TFs and existing gene regulatory networks of arthropods would be of great interest.

## Materials and Methods

### Beetle rearing

*T. castaneum* beetles were cultivated in whole-wheat flour. For sterilization, the flour was kept for 24h at -20°C and another 24h at 50°C. The beetles were maintained inside plastic boxes of approximately 15 × 15cm with humidity between 40-80%.

### Bioinformatic analyses

Protein sequence analyses were performed using BLAST and PFAM searches against proteins available on Genbank and Vectorbase. Dm-zld (NP_608356) was used as initial query for sequence searches. Genomic data from the following genomes were obtained from the Baylor College of Medicine Human Genome Sequencing Center: *Eurytemora affinis, Hyalella azteca, Blattella germanica, Catajapyx aquilonaris, Machilis hrabei, Libellula fulva* and *Ephemera danica*. BLAST results and domain architectures were manually inspected. *In silico* searches in the *T. castaneum* genome for over-represented motifs were performed using the DREME software [61], part of the MEME suite software toolkit [62]. The motif with highest identity with *D. melanogaster zld* binding site (CAGGTAY) was compared to previously described motifs in the FlyFactorSurvey database via TOMTOM. Further, this motif was used to scan 2000 bp upstream of all predicted *T. castaneum* genes with FIMO [14,63,64]. Upstream regions and orthology information of were retrieved from Ensembl Metazoa Biomart (http://metazoa.ensembl.org/biomart/) [65].

### dsRNA synthesis, parental RNAi and embryonic phenotypic analysis

Two non-overlapping PCR fragments containing T7 promoter initiation sites at both ends were used as templates for Ambion T7 Megascript Kit (Cat. No. AM1334) following the manufacturer instructions (for details see Figure S4). The amount and integrity of the dsRNA samples were measured by spectrophotometry and agarose gel electrophoresis, respectively. For parental RNAi (pRNAi) analysis, about 0,5μl of dsRNA was injected from a solution containing 1μg/μl of dsRNA into adult female beetles [35]. Eggs were collected for four egg lays (2 day each) and *zld’s* down regulation was estimated by quantitative Real Time PCR (see below).

### *T. castaneum* embryonic dsRNA injections

Egg injections were performed as previously described [37,42]. Briefly, for the analysis of *Tc-zld* zygotic role, embryos containing nuclear-localized green fluorescent protein (GFP) were collected for one hour and let to develop for additional three hours (30°C) [19,34]. After this period, twenty embryos were dechorionated with bleach (2% solution), aligned into a glass slide and covered with Halocarbon oil 700 (Sigma). Embryos were immediately microinjected at the anterior region with *zld* or *neo* dsRNA at 1 μg/μL concentration with the help of a Nanoinject II instrument (Drummond Scientific Company). After injection, a single nGFP embryo was photographed every five minutes during the following 16 hours (25°C) in a Leica DMI4000 inverted microscope using a GFP filter. Single photographs were used to generate a movie using Windows Movie Maker (supplemental movie 1 and 2). Phenotypes of all injected embryos (*neo* or *zld* dsRNA) were scored at the end of the experiment.

### *T. castaneum* larval dsRNA injections

Larvae were injected with *zld* or *neo* dsRNA as previously described [47]. Knockdown phenotypes in pupae and adult beetles were generated by injection of zld or neo dsRNA solutions at a concentration of 1 μg/μL in the dorsal abdomen of individuals on third and sixth larval instars (n=40). Following injection, larvae were reared in flour at 30°C and collected periodically for RNA extraction and phenotype annotation. Adult beetles were then fixed in ethanol 95% overnight for further morphological analysis.

### Morphological analysis of imaginal disc derived tissues

Antennae, legs, elytra and wings were dissected using forceps and placed in a petri dish for observation. Phenotypic analyses and documentation were performed under a Leica stereoscope model M205.

### PD Index

The methods for wing and elytra measurement and PD index were performed according to described by [49]. Leica AF Lite software was used for the wing measurements. Image properties were adjusted in Adobe Photoshop CS4.

### Quantitative real-time PCR

For experiments using embryos, total RNA was isolated from 100mg of eggs collected from specific development stages (0-3, 3-6 and 6-9 hours after egg laying), ovary and carcass (whole beetle without ovary) using Trizol^®^ (Invitrogen), according to the manufacturer’s instructions. Three independent biological replicates were used for each assay. First strand complementary DNA (cDNA) was synthesized from 2 μg of RNA using Superscript III reverse transcriptase (Invitrogen) and oligo(dT). The cDNA was used as template for real time (RT) PCR analysis using SYBR green based detection. RT-PCR reactions were carried out in triplicate, and melting curves were examined to ensure single products. Results were quantified using the “delta-delta Ct” method and normalized to *rps3* transcript levels, as previously described [66]. Primer sequences used during the study are provided at the supplemental data (Table S5).

### *zld* pRNAi in the hemiptera *Rhodnius prolixus (Rp)*

*zld* cDNA sequence was initially identified by BLAST and included in Figure 1. Parental RNAi against *Rp-zld* was performed as previously described [44].

## Acknowledgements

This work has been supported by Fundação Carlos Chagas Filho de Amparo à Pesquisa do Estado do Rio de Janeiro (FAPERJ) and Conselho Nacional de Desenvolvimento Científico e Tecnológico (CNPq). LA is funded by the Intramural Research Program of National Library of Medicine at the National Institutes of Health, USA. RNdaF and TMV are recipients of the Young Scientist of Rio de Janeiro award (FAPERJ, Brazil). RNdaF would like to thank Prof. Michalis Averof for providing the nuclear GFP line and Prof. Siegfried Roth (Univ. Cologne, Germany) for the support during the beginning of this project.

## Table legends

**Table S1** - Zelda homologs in several Pancrustacea species.

**Table S2** - Gene IDs of 3250 genes with the motif GTAGGTAY in the 2kb upstream regions of *T. castaneum genes* protein-coding genes.

**Table S3** - List of 141 Dm-zld targets during the MZT with one-to-one orthologs among *Tc-zld* predicted targets.

**Table S4** - DAVID functional analysis and enriched ontology terms of the genes listed in Table S3.

**Table S5** - Primers used in quantitative RT-PCR experiments.

## Supplementary material legends

**Figure S1 - The onset of Maternal Zygotic Transition (MZT) in *Tribolium castaneum***. (A-C) Nuclear DAPI staining of *Tribolium* embryos between 0-1 hours (A), 3-6 hours (B) and 6-9 hours after oviposition (C). (D,H) Western-blots of embryonic extracts from 0-1, 0-3, 3-6 and 6-9 hours after oviposition using an antibody against the transcriptionally active form of RNA pol II as previously described (Nestorov et al., 2013a) (D) or an antibody against the α-tubulin protein as a loading control (H). (E,F,G) Immunostaining shows nuclear RNA pol II at 3-6 hours (F) and 6-9 hours (G) but not at 0-1 hour (E) after oviposition. Coupling RNA polymerase II staining with nuclear DAPI staining shows that *Tribolium* zygotic transcription starts between 3-6 hours after egg laying, when the energids reaches the periphery (Figure 2A-C-DAPI and E-G-RNA pol II). Similar results were obtained by western-blots using the same antibody (D,H).

**Figure S2: *Tc-zelda* expression during later developmental stages.** (A) Approx. 13 hours after egg lay (AEL) during the beginning of germ band extension *Tc-zld* is expressed at the posterior growth zone (GZ-arrow). (B) 18-21AEL *zld* is still expressed at the posterior most region (arrow), head lobes (arrowheads) and a single gnathal segment (asterisk). (C) Approx. 39–48 AEL, during early dorsal closure, expression is observed at the nervous system (arrow).

**Figure S3: Schematic drawings of *Tc-zelda* non-overlapping dsRNA constructs.** *Tc-zld* gene corresponds to the Beetlebase ID: TC014798. The following primer pairs were used:. C1-Forward ggccgcggAGCGCATCTTCTCCCTATCA and C1-Reverse cccggggcGCCGTTTTGTCGTTTCTCAT. This primer pair amplifies leads to the amplification of a fragment of 528 bp (349-857). A second primer pair C2-Forward ggccgcggACGACGAGTACCGCTTGACT and C2-Reverse – cccggggcCTTACCACAGGTGTCGCAGA was also used and lead to similar results. This primer pair amplifies a fragment of 476 bp (1823-2279) covering the four zinc-finger domains. The lowercase letters contain the primer sequence used as a template for a second PCR using universal primers which adds a T7 promoter at both sides of the PCR template as previously described [69].

**Figure S4: *Tc-zelda* pRNAi embryos affect early and late patterning.** (A,B) *Tc-twi*, a dorsoventral gene (Handel et al., 2005) acquires an irregular shape at the anterior region during early blastodermal stages (n=2/20) after *zld* RNAi when compared to the control (*dsneo*). (C-F) Early (C,D) and late (E,F) expression of the gap gene *Tc-mlpt* (Savard et al., 2006) in control (C,E) and *zld* RNAi (D,F). *Tc-mlpt* loses its anterior expression domain in *zld* RNAi when compared to control. (G,H) Expression of the segmentation gene *Tc-eve* (Brown et al., 1994) in control (G) and *zld* RNAi (H). *Tc-eve* expression is essential for segmentation via a pair rule circuit [38]. After *zld* RNAi the characteristic *Tc-eve* stripe expression in the growth zone is lost.

**Figure S5: *zld* is required for the generation of the posterior region in the hemiptera *Rhodnius prolixus***. (A,C,E) *R. prolixus* control (*dsneo* embryos). (B,D,F,F’) A representative embryo collected from *R. prolixus zld* pRNAi. (F,F’) Embryo inside the chorion, ventral and dorsal views. (C,E) Embryo removed from the egg shell. (G,H) Schematic drawings of control and *zld* pRNAi embryos. (A) DAPI stainings of control (A,E) and *zld* RNAi (B) embryos. (D) Asterisk denotes an eye which can be observed at the ventral side (F). (D) After dissection the eye can be identified due to its characteristic red pigmentation and shape.

